# Human and Rodent Seizures Demonstrate a Dynamic Interplay with Spreading Depolarizations

**DOI:** 10.1101/2025.01.16.633405

**Authors:** Jacob H. Norby, Daniel Hummel, Noah Ricks, John D. Rolston, Shervin Rahimpour, Juha Voipio, Andrew J. Trevelyan, Elliot H. Smith, R. Ryley Parrish

**Affiliations:** Department of Cell Biology and Physiology, Brigham Young University, Provo, Utah. 84602, USA; Neuroscience Center, Brigham Young University, Provo, UT 84602, USA; Harvard Medical School, Boston, MA 02115, USA; Department of Neurosurgery, University of Utah, Salt Lake City, UT 84112, USA; Faculty of Biological and Environmental Sciences, Molecular and Integrative Biosciences, University of Helsinki, Helsinki, 00014, Finland; Newcastle University Biosciences Institute, Medical School, Framlington Place, Newcastle upon Tyne, NE2 4HH, UK

**Author notes:** **Corresponding authors:** Elliot H. Smith and R. Ryley Parrish.

**Keywords:** Infraslow oscillations, epilepsy, ictal discharge, spreading depression

## Abstract

Seizure termination has been linked to spreading depolarizations (SDs) in experimental epilepsy models, and SDs have recently been suggested to protect against seizures. The precise mechanism, however, remains unclear. Additionally, the co-occurrence of SDs with human seizures remains debated.

In this study, we found that SDs are a prominent feature following ictal events in both human clinical recordings (n=20 seizures from 7 epilepsy patients) using direct-current amplifiers and in the 0 Mg^2+^ model of ictogenesis from rodent brain slices (n=17).

Approximately one-third of rodent seizure-like events (SLEs) were associated with SDs, while all human seizures analyzed had associated infraslow shifts, a hallmark of SDs. SDs were more prominent in the lateral frontal, medial frontal, and lateral temporal lobes, as well as the insular cortex in human patients, but were observed in all recorded brain regions. In rodents, SDs clustered toward the end of ictal events, resulting in significantly shorter SLEs (SLE without SD: 32.60 ± 5.31 s; SLE with SD: 16.03 ± 4.45 s) and delayed onset of subsequent SLEs. These SDs also caused significant AC band silencing compared to SLEs without SDs. Interestingly, SLEs with SDs displayed significantly more low gamma activity during ictal events. Using ion-selective microelectrodes, we found no significant correlation between extracellular [K^+^] levels and SLEs ending in SDs, questioning the role of [K^+^]_o_ in SD induction during seizures.

We observed more heterogeneity in human seizures than in rodent SLEs, with some human seizures showing SD-associated termination and others demonstrating SDs in the middle of ictal events; such intra-seizure SDs were never observed in our rodent model. The human data, collected from patients with intractable epilepsy, demonstrate clear SD propagation during seizures and show that SDs appear and propagate, in multiple brain regions simultaneously with ictal events.

Collectively, these results indicate that SDs are a hallmark of ictal activity and may contribute to seizure termination in both experimental and clinical settings. Furthermore, these findings provide unique insight into the neuronal dynamics that promote SD induction by showing that increased low gamma activity during SLEs is more predictive of SD induction than extracellular [K^+^] levels. We also add further support for the hypothesis that SDs are both anti-seizure and anti-ictogenic as they not only limit and delay subsequent ictal activity but also reduce the duration of SLEs. Taken together, these findings provide rationale for further exploration of SDs as a means to prematurely terminate life-threatening seizures.

## Introduction

The overwhelming majority of seizures terminate naturally, without medical intervention. The process underlying seizure self-termination remains a profound mystery. Understanding this process has obvious utility for shortening and preventing seizures as well as providing therapeutic avenues for the rare seizures that do not self-terminate and can become life-threatening. Many proposed hypotheses about endogenous mechanisms of seizure termination exist,^1,2^ such as increases in adenosine,^3,4^ synaptic vesicle depletion,^5^ reduced input from the seizure source,^6,7^ and energy deficiency.^8,9^ While targeting the adenosine system may be tractable,^10,11^ targeting these other proposed mechanisms are either technologically intractable, or would result in prohibitively deleterious side effects. Investigating other endogenous mechanisms, such as spreading depolarizations (SDs), can open additional strategic paths to terminate seizures prematurely. Along these lines, a recent study has suggested that the mechanism behind seizure termination with electroconvulsive therapy is in fact SDs.^12^

Recently, spreading depressions have received attention as an innate anti-seizure mechanism.^13–15^ While spreading depressions were first discovered in the 1940s during experiments to induce seizures,^16^ their relationship to seizures received limited attention until a study demonstrated their strong association with seizure termination.^17^ Spreading depressions are a consequence of spreading depolarization, a large wave of cellular depolarization that results in neurons experiencing a near-complete loss of membrane resistance.^18,19^ This period of very low membrane resistance is likely due to the opening of almost all membrane ion channels, resulting in the period of spreading depression that often follows the SD and consequently appears to be able to terminate seizures. In addition, tissue “pretreatment” with a SD has been demonstrated to delay the onset of seizures in preclinical models of epilepsy.^14^

Despite all this preclinical data, the relationship between SDs and ictal events remains poorly understood. While SDs occasionally associate with seizure termination, it remains unclear if seizure-associated SDs affect seizure properties. Moreover, the prevalence and mechanisms of induction for seizure-associated SDs are unknown, especially in humans where direct-current recordings are rare. However, high levels of extracellular potassium have been routinely suggested to induce SDs during or following seizures.^20–23^ To address some of these missing links, we aimed in this study to gain direct insight into the prevalence of SDs in both a preclinical model of seizures and in human patients, further exploring the association between potassium dynamics during seizures and the prevalence of SDs.

Our collective results demonstrate that SDs have a significant impact on ictal activity and could contribute to seizures terminate in both experimental and human seizures. Furthermore, propagating SDs are a clear hallmark of seizures from human patients with intractable epilepsy that could impact seizure dynamics, suggesting SDs require increased attention and consideration while monitoring patients with epilepsy. These results also add to our understanding of the mechanisms involved in spontaneous seizure termination and could suggest ways to aid in terminating life-threatening seizures like status epilepticus.

## Materials and Methods

### Ethical approval

Rodents were treated in strict accordance either with the Brigham Young University Animal Research Committee (IACUC) guidelines, which incorporate and exceed current NIH guidelines or with the guidelines of the Home Office United Kingdom and Animals (Scientific Procedures) Act 1986 and approved by the Animal Welfare and Ethical Review Body at Newcastle University. The University of Utah Institutional Review Board approved the human studies, and all participants provided informed consent prior to having their electrophysiological data analyzed.

### Slice preparation

We used WT C57BL/6 (Jackson Laboratory stock no. 000664) mice for all experiments in this study. Experiments were performed on mice 4-12 weeks of age, of both sexes. Mice were housed in individually ventilated cages in a 12 h light–12 h dark lighting regime. Animals received food and water *ad libitum*. Mice were sacrificed by cervical dislocation, brains were removed and then placed in cold cutting solution perfused with carbogen (95% O2 and 5% CO2) and containing the following (in mM): 3 MgCl2; 126 NaCl; 26 NaHCO3; 3.5 KCl; 1.26 NaH2PO4; 10 glucose. 400 µm horizontal sections of brains were made for all extracellular recordings while 350 µm coronal sections were made for all patch-clamp recordings, on a Leica VT1200 vibratome (Leica Microsystem). Slices were stored at room temperature, in a holding chamber for 1-4 h before experimentation. Holding chamber solutions were bubbled with carbogen (95% O2 and 5% CO2) in aCSF containing the following (in mM): 2 CaCl2; 1 MgCl2; 126 NaCl; 26 NaHCO3; 3.5 KCl; 1.26 NaH2PO4; 10 glucose.

### Extracellular recordings

Recordings were performed using an interface recording chamber. Slices were placed in the recording chamber and perfused with the same aCSF solution as used in the holding chamber expect for the removal of the Mg^2+^ ions to induce ictal-like activity. Recordings were obtained 1-3 MΩ borosilicate glass microelectrodes (GC120TF-10; Harvard using aCSF-filled apparatus) placed in deep layers of neocortex. Extracellular potassium, [K ]_o_, was measured using single-barreled K^+^-selective microelectrodes. The pipettes were pulled from nonfilamented borosilicate glass (Harvard Apparatus), and the glass was exposed to vapor of dimethyl-trimethyl-silylamine (Sigma-Aldrich), baking at 200°C for 40 min. The pipettes were then backfilled with aCSF. A short column of the K^+^ sensor (Potassium ionophore I, cocktail B; Sigma-Aldrich, #99373) was taken into the tip of the salinized pipette by using slight suction.

The recordings through the K^+^-sensor electrode were referenced to a second electrode filled with aCSF. From the differential signal from a custom build amplifier, we calculated the [K^+^]_o_ from calibration recordings made in an open bath, using sudden increments in [K^+^]_o_. We checked the stability of the electrodes at the start and end of each recording. Data from unstable electrode recordings were discarded. This provided a scaling factor S, of 55-59 mV, where the K^+^ concentration at a given moment in time, t, was calculated from the differential voltage, V(t), as follows: [K^+^]=[K^+^]_baseline_10^V(t)/S^. [K^+^]_baseline_ for our experiments was 3.5 mM. The temperature of the chamber and perfusate was maintained at 33°C–36°C using a closed circulating heater (FH16D, Grant Instruments). The solutions were perfused at the rate of 3 ml/min by a Watson Marlow 501U peristaltic pump (Watson-Marlow Pumps). The DC-coupled local field potential (LFP) signal was unfiltered and amplified to a 10× output with a custom build amplifier, while the alternating current LFP signal had a bandpass filter (1-3000 Hz), amplified (gain:100) using a Digitimer NeuroLog System (NL106 AC/DC Amplifier). These waveform signals were digitized with a Micro 1401-3 ADC board (Cambridge Electronic DesignK) and Spike2 version 7.10 software (Cambridge Electronic Design). Signals were sampled at 10 kHz. Recordings were analyzed using a custom-written code in MATLAB (The MathWorks, Natick, MA, USA).

### Patch-clamp electrophysiology

Recordings were made using a Multiclamp 700B amplifier (Molecular Devices, San Jose, CA, USA) and pClamp software, digitized at 10 kHz. The bath was mounted on a Scientifica movable top plate (Scientifica, Uckfield, UK) fitted with a heater plate (Warner Instruments, Hamden, CT, USA), and the incoming solution (perfusion at 2–3 ml/min) was heated by a sleeve heater element (Warner Instruments). All imaging and electrophysiological recordings were done at 33–37°C. Whole-cell patch-clamp recordings of deep-layer neocortex pyramidal cells were made using 4–7 MΩ pipettes (borosilicate glass; Harvard Apparatus, Camborne, UK) controlled with PatchStar micromanipulators (Scientifica). Pipettes were filled with a KMeSO4-based internal solution containing (in mM): 125 KMeSO4, 6 NaCl, 10 Hepes, 2.5 Mg-ATP, 0.3 Na2- GTP, 0.5% (w/v) biocytin, 5 mM QX-314 (N-(2,6-dimethylphenylcarbamoylmethyl) triethylammoniumbromide). All experiments were run in ACSF without Mg^2+^ to induce ictal-like activity. The electrophysiological data were analyzed off-line using in-house software implemented in MATLAB (The MathWorks, Natick, MA, USA).

### Viral injections

WT C57BL/6 pups were injected with AAV9.Syn.flex.GCaMP6f, purchased from the University of Pennsylvania vector core. Injections were performed either on postnatal day 0 or 1. Pups were set on a stereotaxic frame and anaesthetized and maintained with 2% isoflurane, following application of EMLA cream (2.5% lidocaine and 2.5% prilocaine) to the left top of their head. Injections were made, using 10 μl Hamilton syringes with a beveled 36-gauge needle World Precision Instruments, Sarasota, F USA), unilaterally into the lateral ventricle and overlying cortical plate, at about 1 mm anterior to lambda and 1 mm lateral to the midline into the left hemisphere, starting at 1.7 mm depth from the top of the dura mater, for a total of four separate 50 nl injections, deepest first and coming up 0.3 mm for each subsequent injection. Approximately 200 nl (~1000 viral particles) were injected into the left hemisphere over a 10 min period. Pups were m∼onitored until they awoke following the procedure and then returned to their home cage. These neonatal injections produced widespread cortical expression of GCaMP6f into the neurons of interest.

### Live tissue imaging

Live imaging experiments were performed using a spinning disc confocal microscope (UMPlanFL N ×20, 0.5 NA objective). The tissue was illuminated with a 491 nm laser (Cobolt Calypso 50; Cobolt AB, Solna, Sweden) for visualization of the GCaMP6f. Imaging was made using a Hamamatsu C9100 EM camera (Hamamatsu Photonics (UK)), Welwyn Garden City, UK), run by VoxCell software (Visitech International, Sunderland, UK; microscope) on Dell Precision computers (Dell Technologies, Round Rock, TX, USA). Imaging was performed at 10 Hz. Off-line analysis of the images was performed using ImageJ (imagej.net, NIH, Bethesda, MD, USA) and in-house software implemented in MATLAB (The MathWorks, Natick, MA, USA).

### Human data methods

Human participants were undergoing neuromonitoring for surgical treatment of their drug-resistant epilepsy. In each participant, electrophysiological recordings of seizures were obtained from intracranially implanted electrodes: stereo-electroencephalographic (sEEG) depth electrodes and electrocorticographic (ECoG) surface electrodes. Seizures, in which there was sufficient spatial coverage of infraslow shift migration, from each participant were trimmed from the clinical record and analyzed in three frequency bands: direct-current (DC), local field potential (LFP), and phase-locked high gamma (PLHG). DC signals were filtered in matlab using a non-causal, 96th-order finite impulse response (FIR) stopband filter with a cutoff frequency of 0.05 Hz relative to the new sampling frequency. The signals were first resampled to 10 Hz and then processed using the filtfilt function to ensure zero-phase distortion. LFP signals were filtered above 0.5 Hz using a non-causal, 4th order Butterworth filter in matlab. PLHG was calculated from Morlet wavelet decompositions of the signal on each channel, where the high frequency power between 80 and 150 Hz was weighted by the low frequency phase between 1 and 25 Hz.24

Putative SDs were identified by an infraslow shift of a magnitude greater than 2 mV and seizures by a PLHG strength above 0.01 mV. To assess whether these putative SDs propagated, 0.05 Hz high-pass filtered LFP traces from adjacent channels (on the same recording shank) to identified SD events were extracted for analysis. The time region of analysis was from the beginning of the first identified seizure innervation (as determined by an above threshold PLHG value) until approximately 5 minutes after seizure termination. Electrodes that did not achieve an infraslow shift with a magnitude greater than 2 mV were considered not recruited by the SD and not included in the propagation analysis. The propagation of seizure-associated infraslow shifts throughout the brain was measured using the minimum value for each infraslow LFP trace in and adjacent to seizure and SD recruited areas. The electrode to experience an infraslow minimum of greater magnitude than 2 mV at the earliest time point was denoted as the putative SD induction site, from which the distances of other electrodes were then calculated. The time of each minimum and the distance from the putative SD induction site were linearly regressed and the regression validated with a permutation test. This analysis method was also repeated for maxima and extrema to determine how propagation might be observed with these features (see Supplemental Fig. 3).

To characterize the spatial aspects for the implanted DC electrodes in human patients, we used the LeGUI software^25^ to identify brain area localization based on post-implant CT and preoperative MRI scans through coregistration and normalization to the Montreal Neurological Institute (MNI) template. LeGUI Electrode detection is automated for both surface and depth electrodes, and image processing capabilities are powered by the MATLAB Statistical Parametric Mapping toolbox.

### Statistical analyses

Statistical analysis of data was performed using custom MATLAB code and automated where possible. Data were analyzed with two-tailed t-tests for all bar plots of two groups (Fig. 1, Fig. 2, Fig.3, Fig 4, Supplemental Fig. 1). Data were also analyzed with Chi-squared McNemar tests (Table 1) and mixed-effects derived ANOVA models and permutation tests (Fig. 6, Supplemental Fig. 3). Permutation tests were performed by shuffling the location and time pairings of data for *n* permutations and calculating the ANOVA *F* statistic (with mixed effects model if data from multiple patients were present: [Delay ∼ Distance + (Distance|Patient)]) for each permutation. Pairwise post-hoc chi-square tests with Bonferroni correction for multiple comparisons and G-tests were used to analyze data in supplemental analyses (Supplemental Fig. 2). Statistical significance was determined using *⍰*<0.05, except with tests using the Bonferroni correction. For voltage clamp data, the time when pre-ictal activity was determined to return was when the first spike of comparable amplitude to pre-ictal spikes occurred. The [K^+^]_o_ rate of rise was determined by taking the slope of the linear regression of the [K^+^]_o_ trace during the rising phase of the SLE. The area-under-the-curve (AUC) was calculated using the trapz function. [K^+^]_o_ analyses were performed using custom MATLAB code. Figures were made in MATLAB (The MathWorks, Natick, MA, USA).

**Figure 1.**
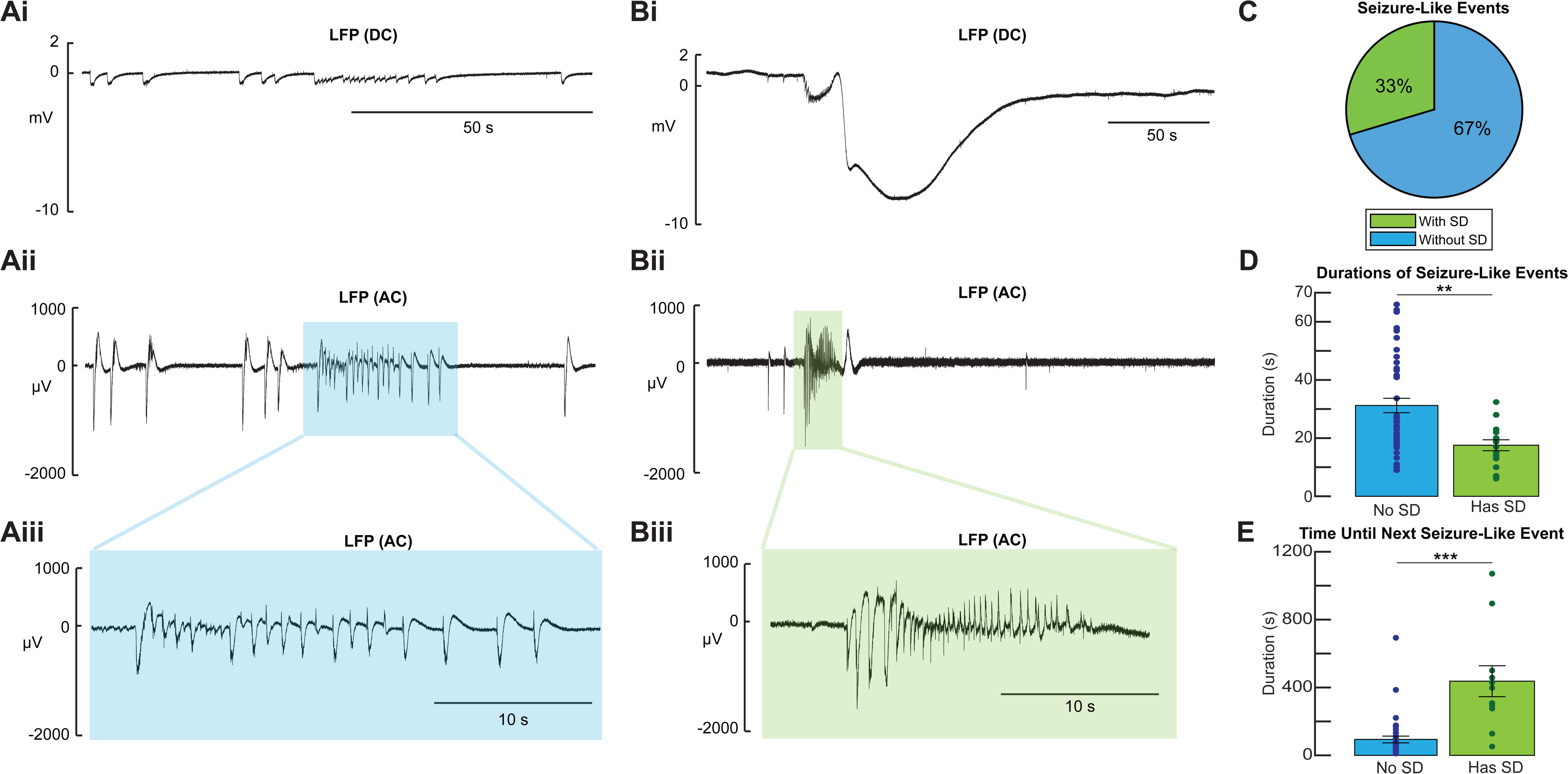
SDs decrease the duration of SLEs and increase the time to subsequent SLEs. Representative LFP traces of **(A)** a SLE terminating without a SD and **(B)** a SLE terminating with a SD, including DC **[i]** and AC **[ii]** traces, as well as **[iii]** zoomed in traces of the SLEs. **(C)** Quantification of the proportions of SLEs that terminated in a SD and those that did not. **(D)** SLEs that ended in SDs were significantly shorter than seizures that did not end in SDs (SLE w/o SD; n= 41, SLE with SD; n= 15, two-tailed t-test, p= 0.0028). **(E)** The time to next SLE following the termination of a SLE with a SD was significantly greater than following a SLE without a SD (SLE w/o SD; n= 37, SLE with SD; n= 11; two-tailed t-test, p= 1.3 x 10^-6^). All error bars represent SEM.

**Figure 2.**
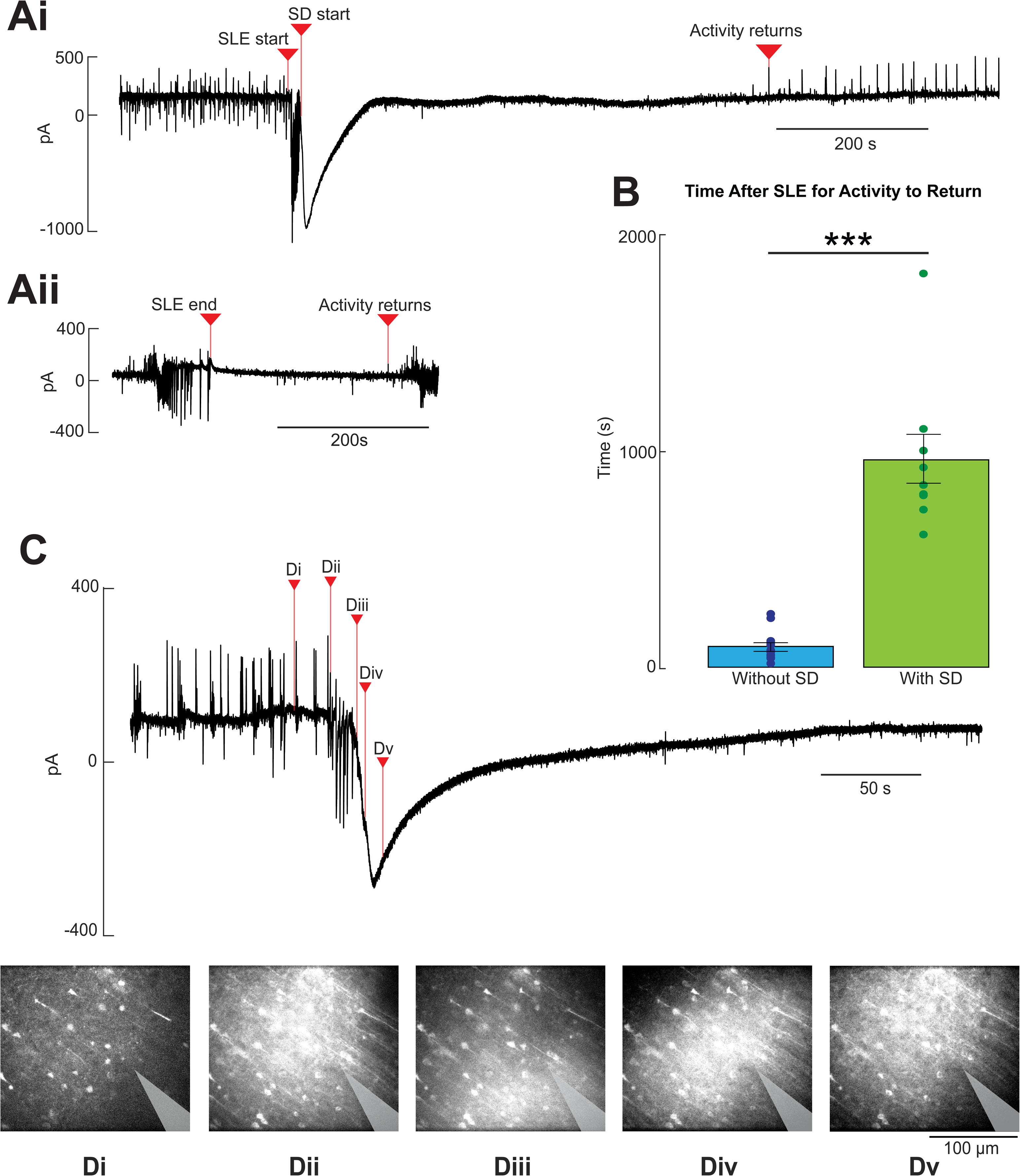
SDs delay the return of neuronal activity following SLEs. **(A)** Representative patch- clamp recordings highlighting a SLE terminating in a SD **[Ai]** and a SLE that terminates without a SD **[Aii**]. **(B)** SLEs terminating with a SD are followed by significantly longer durations of network quiescence compared to SLEs that did not terminate in a SD (SLEs w/o SD, n= 16, SLEs with SD, n=9; two-tailed t-test, p= 1.5 x 10^-9^). Error bars represent SEM. **(C)** Representative patch-clamp trace of a SLE terminating in a SD while Ca2+were imaged in the neurons. **(D)** Selected frames from Ca2+ imaging from before the start of the SLE **[Di]**, during a SLE **[Dii]**, and at various points as a SD propagates from the bottom right of the frame diagonally upwards **[D(iii-v)]**.

**Figure 3.**
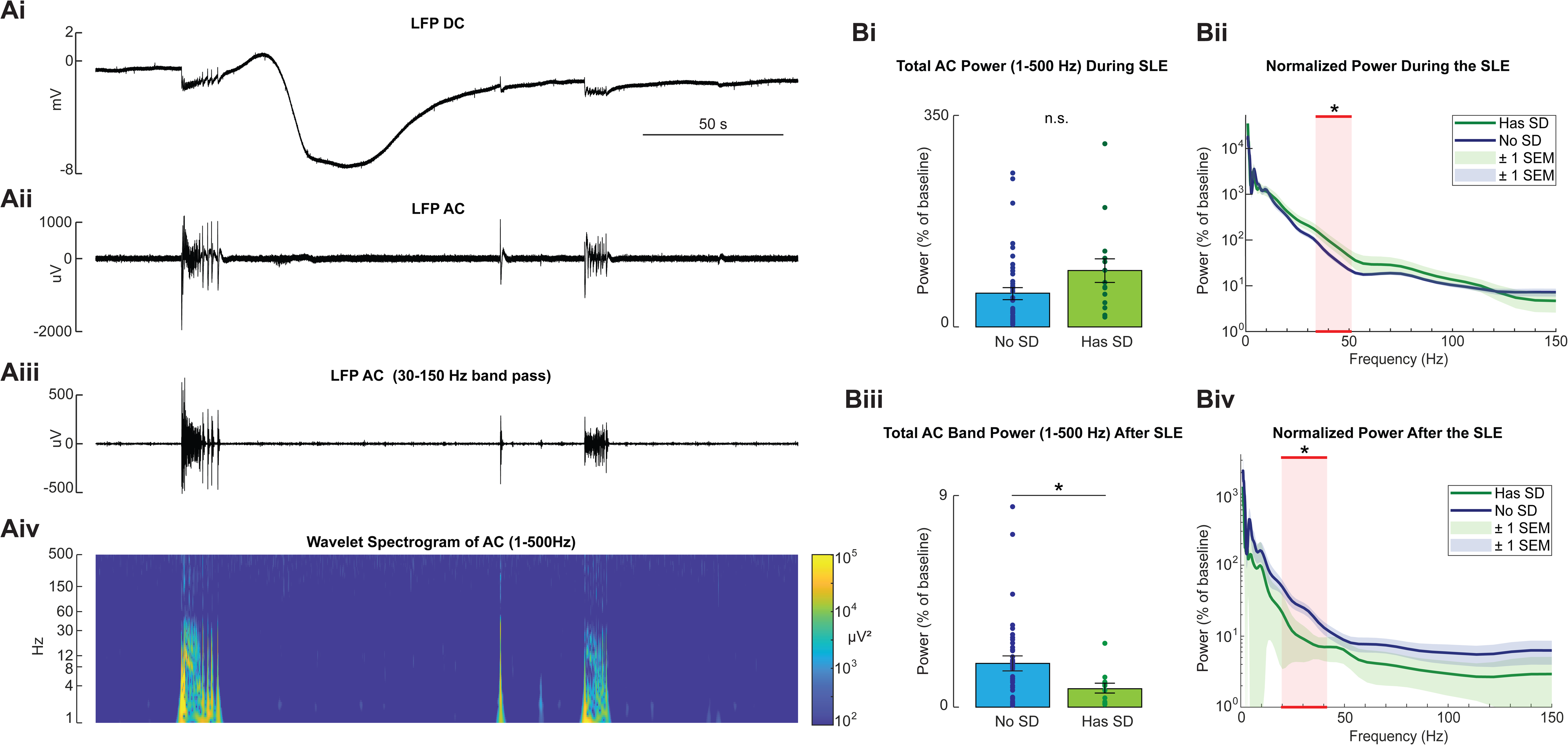
AC low gamma activity is increased during seizures that end in SDs while total AC band activity is reduced following SDs. **(A)** Representative DC LFP trace **[Ai]**, AC LFP trace **[Aii]**, AC LFP trace with a band pass filter from 30-150 Hz (highlighting gamma frequency activity) **[Aiii]**, and wavelet spectrogram of the LFP (AC) trace **[Aiv]** of a SLE terminating in a SD followed by a SLE not terminating in SD. **(B)** The normalized power of spectral activity (given as a percentage of baseline activity, normalized by time) for SLEs terminating in SDs and SLEs without SDs (SLEs w/o SD, n= 41; SLEs with a SD, n= 15). Total AC power (1-500 Hz) during SLEs trended towards greater power for SLEs that terminated in a SD, yet this difference was not significant (two-tailed t-test, p= 0.063) **[Bi]**. AC power in the lower gamma frequencies (30-50 Hz), was significantly higher during SLEs that did terminate in a SD (two-tailed t-test, p= 0.0011) **[Bii]**. Total AC power (1-500 Hz) for 120 s following SLEs was significantly reduced for SLEs that terminate in a SD compared to SLEs without a SD (two-tailed t-test, p= 0.029), indicating total AC band silencing following SLEs that terminate in a SD **[Biii]**. Especially reduced spectral power was observed in the low gamma (30-50 Hz) frequency range (two-tailed t-test, p= 0.0049) for 120 s following SLEs **[Biv**]. All error bars represent SEM.

**Figure 4.**
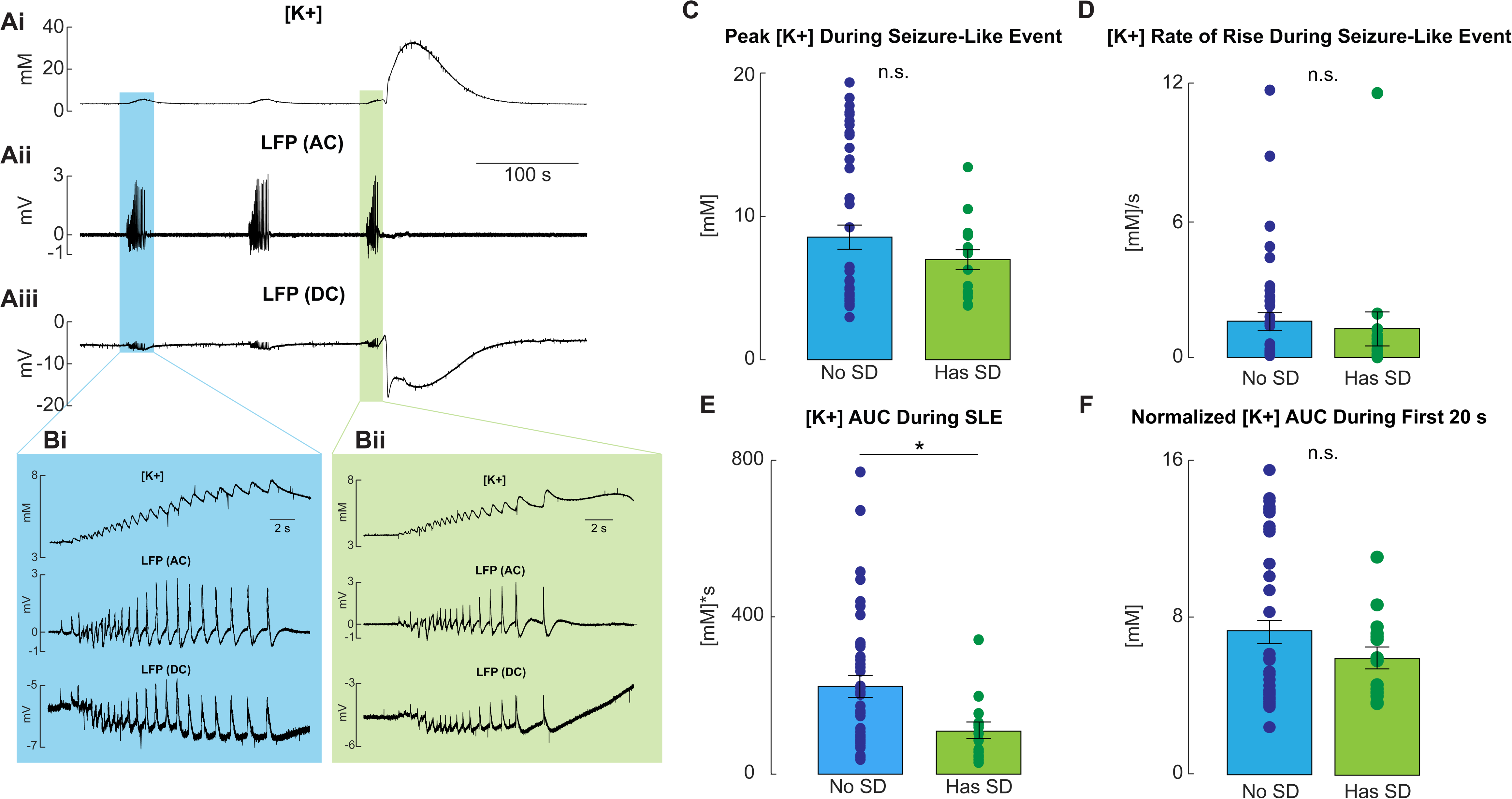
Extracellular [K^+^] concentrations do not correlate with the presentation of SDs during or following SLEs. **(A)** Representative [K^+^] trace **[Ai]**, LFP (AC) trace **[Aii]**, and LFP (DC) trace **[Aiii]**. **(B)** Zoomed-in traces from **A** that highlight a SLE that terminated without a SD **[Bi]** and a SLE that terminates with a SD **[Bii]**. **(C)** Peak [K^+^] reached during a SLE was not found to be significantly different between SLEs that terminated in a SD and those that did not (SLEs w/o SD, n= 41; SLEs with a SD, n= 15; two-tailed t-test, p=0.321). **(D)** [K^+^] rate of rise during a SLE was not found to be significantly different between SLEs that terminated in a SD and those that did not (SLEs w/o SD, n= 41; SLEs with a SD, n= 15; two-tailed t-test, p=0.669). **(E)** The [K^+^] area under the curve during SLEs was found to be significantly greater for SLEs not terminating in a SD and those that did terminate in a SD (two-tailed t-test, p=0.019). **(F)** The [K^+^] area under the curve, normalized to time, during the first 20 s of SLEs was not found to be significantly different between SLEs not terminating in a SD and those that did not terminate in a SD (two-tailed t-test, p=0.223). All error bars represent SEM.

## Results

### Spreading depolarizations (SDs) shorten seizure duration and delay the onset to subsequent seizures

Experimental seizures and SDs have been demonstrated to co-occur in many studies,^15,17,26–30^ however neither the prevalence nor the consequences of SDs have been fully documented. To gain insight into the frequency of SDs and acute network changes induced by SDs in an *ex vivo* seizure model, we used a custom-made head stage split between an AC and DC amplifiers. This set-up allowed us to study infraslow oscillations (below 0.05Hz, where SD occurs) and high frequency activity (greater than 1Hz) indicative of seizures from the same location in rodent neocortex. To induce seizure-like events (SLEs) in rodent brain slices, Mg^2+^ ions were removed from the aCSF (Fig. 1A, B). Using this model of ictogenesis, we observed SLEs terminating both with, and without, SDs; often both occurred in the same brain slice.

Collectively from all our *ex vivo* rodent recordings in this study (LFP and patch-clamp recordings, totaling 17 brain slices and 73 SLE), we found that 33% of the SLEs exhibited SD at seizure termination (Fig. 1C).

It has also been suggested that SDs can bring about seizure termination.^13,15,17,31,32^ Therefore, we next sought to determine if SDs had any impact on SLE properties. To determine this, we analyzed the duration of SLEs with and without an associated SD. We found that when SLEs terminated with an associated SD, the duration of the SLEs was significantly shorter than SLEs that did not associate with SDs (Fig. 1D; *t*(54)= -3.14; *p=* 0.0028). Additionally, the time until subsequent SLEs following SD-terminated SLEs was significantly increased when compared to SLEs that did not present with a SD (Fig. 1E; *t*(46)*=* 5.57; *p=* 1.3 × 10^-6^) as has been previously reported.^14^

In line with this finding, using voltage-clamp recordings, we also found that there was significant silencing of all 0 Mg^2+^ induced epileptiform activity of SLEs following a SD compared to SLEs that did not terminate with a SD (Fig. 2A, B; *t*(23)*=* 9.63; *p*= 1.5 x 10^-9^).

These findings suggest that SDs may be a potential anti-ictogenic mechanism that both curtails and is protective against ictal activity. This termination of a SLE can be visualized by Ca2+ imaging during a simultaneous voltage-clamp recording, demonstrating a SD wave migrating into an “ictal core,” coinciding with the end of an ictal event (Fig. 2C, D; Supplemental Video 1). This collective data is consistent with the hypothesis that SDs can act as an endogenous antiseizure mechanism.

### Ictal events that end in SDs display altered AC spectral features during and after SLEs

It is well known that SDs typically result in significant cortical silencing that can last for many minutes, termed a spreading depression.^14,19,33–35^ This strong period of depression could be a major contributory factor towards suppressing subsequent epileptiform activity. We therefore wanted to explore if there were apparent changes in our AC recordings that suggest a strong depression following the SLEs. However, we first investigated the spectral features during the SLE (Fig. 3A) to determine if there were any changes in the AC bands that might predict a future or already migrating SD wave. During the SLE, we found no overall increase in total AC power (1-500 Hz) (Fig. 3Bi; *t*(54)*=* 1.90; *p=* 0.063), however AC power in the low gamma frequencies (30-50 Hz) was found to be significantly greater during SLEs that were followed by SDs (Fig. 3Bii; *t*(54)*=* 3.45; *p=* 0.0011; SLE w/o SD: *n=* 41, SLE with SD: *n=* 15; two-tailed t-test), suggesting that an observable network change is present during the SLE before the tissue induces a SD or possibly prior to the tissue being innervated by a SD wave.

In addition to this observed increase in low gamma power during the SLE, there was a pronounced silencing effect following SLEs associated with SDs lasting several minutes with a mean duration of 106.2 s and a standard deviation of 30.7 s. SLEs that associated with SDs showed a significantly reduced AC power for 120 s following the termination of the SLE compared to SLEs that did not associate with SDs (Fig. 3Biii; *t*(54)*=* -2.25; *p=* 0.029).

Interestingly, we observed significantly reduced power in the low gamma frequencies following SLEs with SDs (Fig. 3Biv; *t*(54)*=* -2.94; *p=* 0.0049), largely the same frequencies that increased during SLEs with SDs. This data suggests that the commonly observed phenomena of spreading depression following SDs can contribute to a post-ictal decrease in epileptiform activity but is unlikely to be the only cause of the delay of subsequent seizures. This decrease in AC band power only lasts for a few minutes following the SD, however it can take well over 10 minutes for epileptiform activity to return (Fig. 2B).

### Extracellular potassium does not correlate with SD induction during or following ictal discharges

Since high [K^+^]_o_ is a commonly accepted mechanism for SD initiation,^20,36–38^ and SDs can be consistently and reliably induced experimentally via application of extracellular [K^+^],^39–41^ we next sought to test the hypothesis that the well-documented increases in [K^+^]_o_ during SLEs predict SD induction.^42–47^ Using ion-selective microelectrodes, we measured [K^+^]_o_ during both SLEs with and without SDs (Fig 4A, B). Notably, we found no significant differences in peak [K^+^]_o_ during or after SLEs with or without SDs (Fig. 4C; *t*(54)*=* -1.00; *p=* 0.321; see also Supplemental Fig. 1). Multiple SLEs reached [K^+^]_o_ levels as high as 20 mM yet still did not present with an associated SD (Fig. 4C). To determine if the rate of [K^+^]_o_ increase was more predictive of SLE-associated SD, we characterized the rate of [K^+^]_o_ rise during the rising phase of the SLE and found that the rate of [K^+^]_o_ rise was also not correlated with SD incidence (Fig. 4D; *t*(54)*=* -0.429; *p=* 0.669). We next approximated the difference in bulk [K^+^]_o_ change during the SLE by measuring the area-under-the-curve (AUC) of the [K^+^]_o_ trace and found that the total [K^+^]_o_ load of SLEs was significantly higher for SLEs that did not associate with a SD (Fig. 4E; *t*(54)*=* -2.42; *p=* 0.019). This increased [K^+^]_o_ load was primarily driven by the previously shown significant increase of duration for SLEs without SDs (see Fig. 1D). However, this data further demonstrates that SLEs without SDs had a more significant extracellular potassium load than the SLEs with SDs, yet this large, sustained [K^+^]_o_ change never induced a SD. When the [K^+^]_o_ AUC is normalized by the SLE durations, the difference in [K^+^]_o_ load was not significant (Supplemental Fig. 1A; *t*(54)*=* -1.14; *p=* 0.257). To further test for potential obscuring of trends in the [K^+^]_o_ load by measuring across the entire SLE, we additionally tested the normalized (to time) [K^+^]_o_ AUC during the first 20 s of the SLE, as 20 s was approximately the mean time by which SLEs that did terminate in a SD had triggered the SD (see Fig 1D). This test showed no significant connection between the normalized initial 20 s [K^+^]_o_ AUC and if a SLE would terminate in a SD (Fig. 4F; *t*(54)*=* -1.233; *p=* 0.223). Collectively, this data suggests that increases in [K^+^]_o_ caused by seizures is not by itself a key driver of SD induction.

### Seizures and infraslow oscillations co-occur in humans

While SDs are known to be a prominent feature during experimental seizures,^14–16^ their prevalence during human seizures remains debated.^48,49^ While a few studies have investigated how seizures and SDs co-occur in patients with epilepsy,^29,50–52^ how these phenomena may present across various brain regions, and their relationship with seizure activity remains poorly understood. Therefore, using electrophysiological recordings of 20 seizures from clinical electrodes implanted in 7 human patients (mean of 3 ± 1.07 seizures per patient), we sought to characterize the spatiotemporal dynamics of seizure-associated infraslow oscillations (<0.05 Hz) (see Fig. 5A, ^53,54^). Phase-locked high gamma (PLHG) served as our proxy to determine seizure recruitment on each electrode ( Fig. 5B).^55^ We observed infraslow shifts of multiple millivolts in at least some region of the brain (Fig. 5 C, D; Supplemental Video 2). All 20 seizures for the 7 patients exhibited these seizure-associated infraslow shifts in at least one brain area. We also observed that the incidence of these large infraslow shifts sometimes associated with silenced neuronal activity in conjunction with the infraslow shift (Fig. 6 A, B, C). These large infraslow shifts were found to occur both during and after ictal events, demonstrating more diversity than we observed in our rodent data, where SDs primarily occurred following the SLEs. Collectively, these findings suggest that these putative SDs consistently co-occur with seizures in spontaneous human seizures.

**Figure 5.**
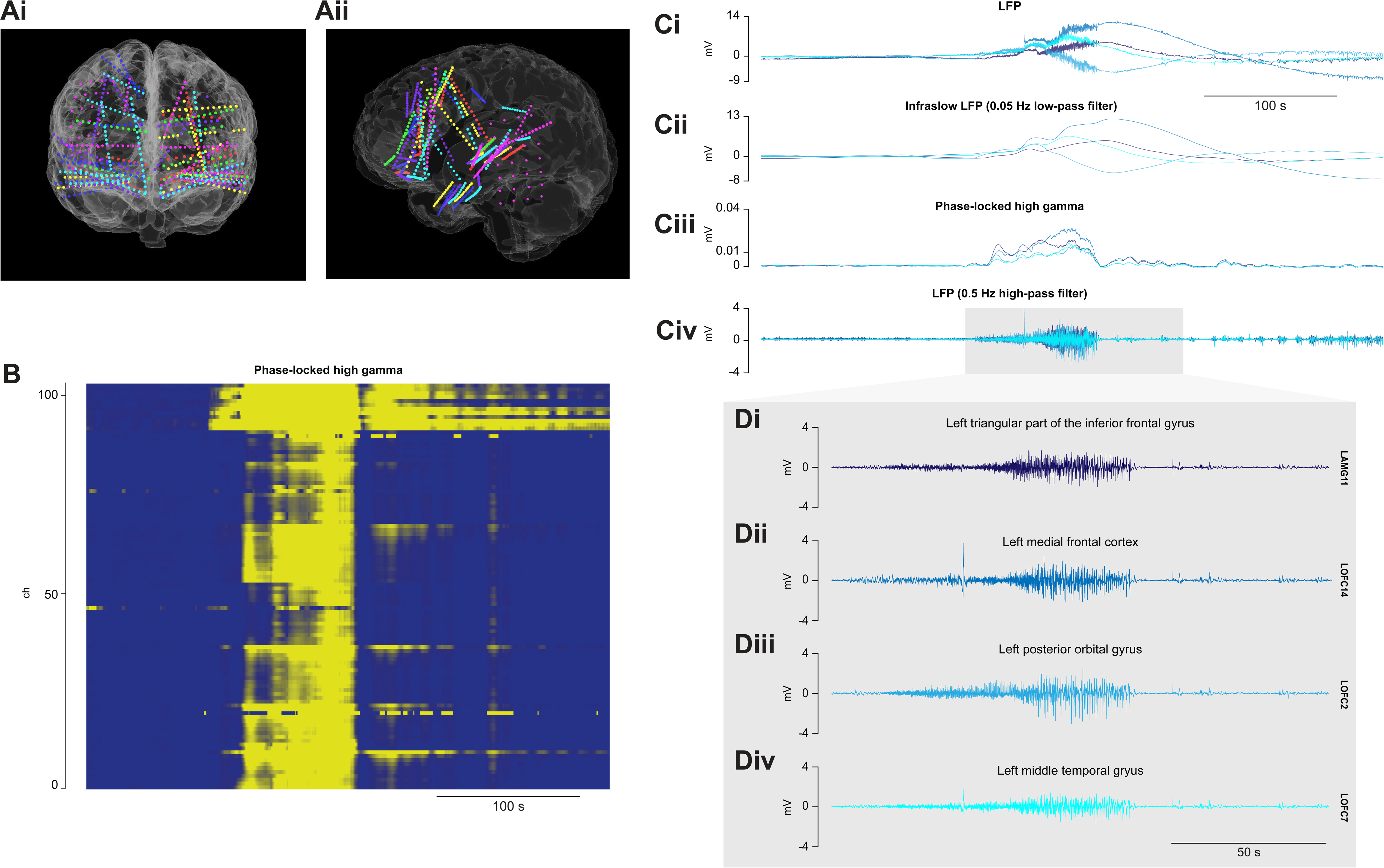
Infraslow oscillations indicative of SDs co-occur with seizures in human patients. Coronal **[Ai]** and sagittal **[Aii]** views of overlaid 3D reconstructions of the patients’ brains (n= 7) based on CT/MRI scans. Each patient’s electrode locations are color-coded on the 3D brain overlay. **(B)** Representative phase-locked high gamma raster plot of all electrodes during a seizure. **(C)** A representative intracranial macroelectrode recording highlighting the seizure from **B** and associated infraslow shift from four recording sites in the frontal and temporal lobes. The broadband LFP signal **[Ci]**, the infraslow LFP signal low-pass filtered below 0.05 Hz **[Cii**], the phase-locked high gamma activity **[Ciii]**, and the AC LFP activity high-pass filtered above 0.5 Hz **[Civ]**. **(D)** Zoomed-in individual LFP (0.5 Hz high-pass filtered) traces from **Civ** for each recording site during the seizure.

**Figure 6.**
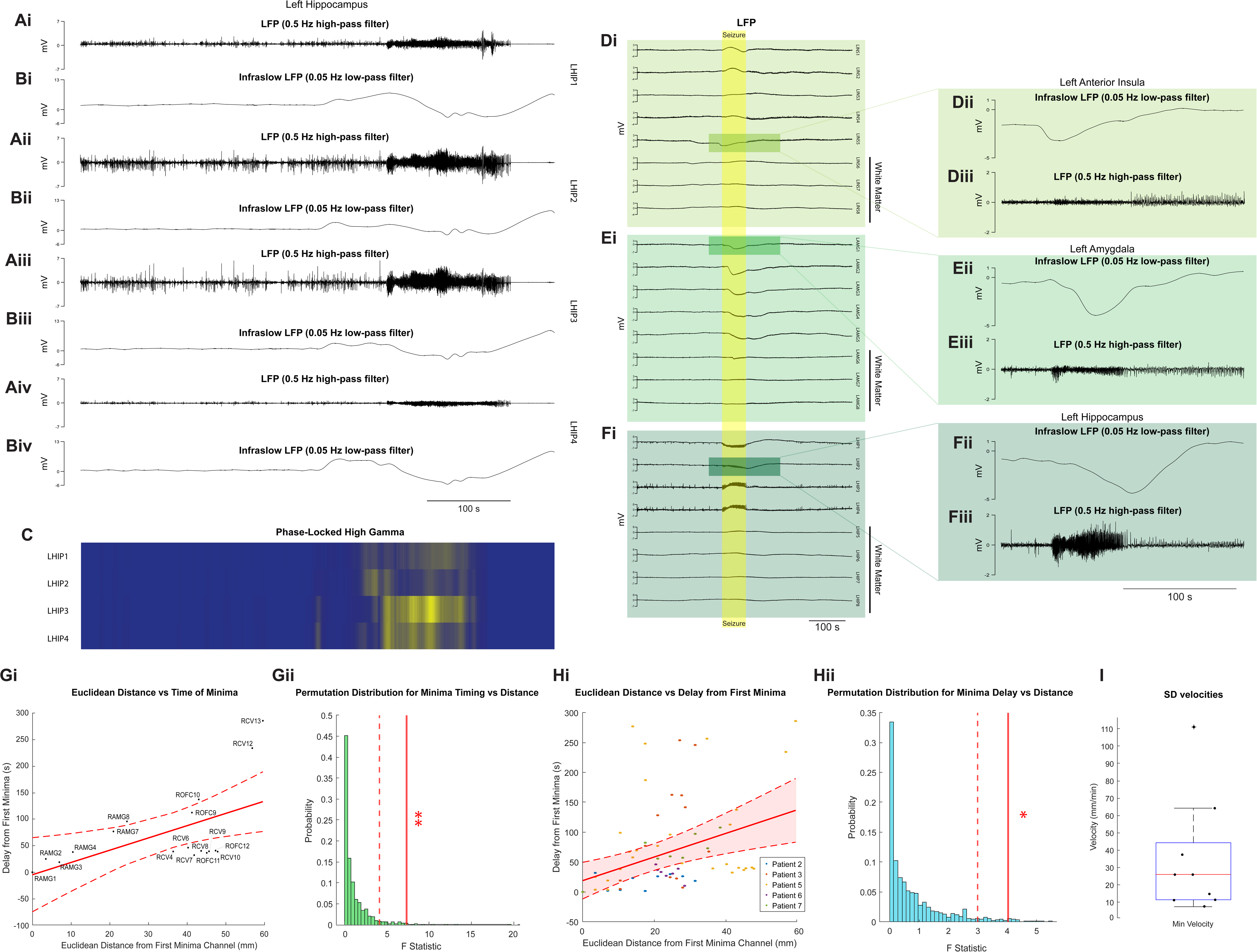
Infraslow waves observed in human patients can associate with seizure termination and propagate similarly to SDs. (A-C) Representative traces of a human seizure demonstrating AC silencing following a seizure that terminated with a co-occurring SD. **(A)** The AC LFP (0.5 Hz high-pass filtered) traces for the electrodes in the left hippocampus **[i-iv]**. **(B)** The infraslow LFP (0.05 Hz low-pass filtered) traces electrodes in the hippocampus **[i-iv]**. **(C)** The phase-locked high gamma associated with the traces shown in **A**. **(D-F**) Representative traces from various brain areas showing propagation of a SD wave which co-occurs with a seizure. **(D)** Representative LFP traces from electrodes (each spaced ∼5 mm apart) implanted in left posterior insula (LINS1-2), left anterior insula (LINS3-5), left frontal operculum (LINS6), and left cerebral white matter (LINS7-8). **[Di]** The broadband LFP trace, **[Dii]** the 0.05 Hz low- pass filtered infraslow LFP , and **[Diii]** the 0.5 Hz high-pass filtered AC LFP. **(E)** The LFP traces from electrodes (each spaced ∼5 mm apart) implanted in the left amygdala (LAMG1-5) and left cerebral white matter (LAMG6-8). **[Ei]** The broadband LFP trace, **[Eii]** the 0.05 Hz low-pass filtered infraslow LFP and **[Eiii]** the 0.5 Hz high-pass filtered AC LFP. **(F)** The LFP traces from electrodes (each spaced ∼5 mm apart) implanted in left hippocampus (LHIP1-5) and left cerebral white matter (LHIP5-8). **[Fi]** The broadband LFP trace, **[Fii]** the 0.05 Hz low-pass filtered infraslow LFP, and **[Fiii]** the 0.5 Hz high-pass filtered AC LFP**. (G**) Representative propagation of a SD wave. **[Gi]** Correlation of selected electrode shanks showing temporal delay (vertical axis) and distance from the first channel (RAMG1) to display a minimum infraslow shift co- occurring with a seizure. The solid line represents the line of best fit and the dashed lines the 95% confidence interval (*n*= 19 electrodes, *r*= 0.56, velocity= 25.92mm/min). **[Gii]** The permutation distribution plot of ANOVA F-statistics from 1000 permutations shuffling the minima delays of the infraslow shifts and electrode locations (*p*= 0.009). **(H**) Combined propagation of SD waves is significant when propagating SD waves were identified by their minima. **[Hi]** Correlation of electrodes showing temporal delay (vertical axis) and distance from the first infraslow shift minima (of each seizure recording) to display an infraslow shift that co- occurs with a seizure. The solid line represents the line of best fit and the dashed lines the 95% confidence interval (*n*= 103 electrodes from 9 seizures, *r*= 0.52, velocity= 35.49 mm/min). **[Hii]** The permutation distribution plot of the ANOVA F-statistics (derived from the mixed effects model) over 10000 permutations of the minimum infraslow shift delays and electrode locations (*p*= 0.027). **(I)** Distribution of the SD velocities for the 9 seizures from H.

In several brain regions, seizures and SDs also co-localized more often than would be expected by chance. Seizures were associated most commonly with SDs in the lateral and medial frontal lobes, as well as the lateral temporal lobe, insula, hippocampus, and amygdala (Table 1; lateral frontal lobe: χ^2^(1)*=* 44.26, *p=* 2.87 x 10^-11^; medial frontal lobe: 2(1)*=* 28.47, *p=* 9.51 x 10^-8^; lateral temp χ lobe: 2(1)*=* 51.98, *p=* 5.61 x 10^-13^; insula: 2(1) χ 7.20, *p=* 0.0073; oral *=*2 2 hippocampus: χ (1)*=* 6.53, *p=* 0.011; amygdala: χ (1)*=* 6.53, *p=* 0.011; McNemar test). Occipital, parietal, and subcortical areas were only measured in a limited sample of patients and were not found to have significant patterns of co-occurrence (Table 1; occipital: 2(1)*=* 0.67, *p=* 0.41; parietal: 2(1)*=* 3.00, *p=* 0.08; subcortical: 2(1)*=* 0.20, *p=* 0.65; McNemaχtest; see χ χ r Supplemental Fig. 2). Although strong DC shifts indicative of SDs were found in the thalamus, only one electrode was found to be recruited into a seizure and did not co-occur with a SD (Table 1). These findings strongly indicate that seizures and SDs co-occur for human epilepsy patients.

### Infraslow oscillations propagate during and after human seizures

While these large infraslow oscillations have characteristics of SDs such as the large infraslow shift, SDs typically demonstrate a slow propagating wave.^30,34,41,56,57^ We therefore sought to determine whether seizure-associated infraslow oscillations propagated as SDs. The placement of multiple electrode shanks throughout the brain not only allowed us to measure each brain region individually but also permitted us to track seizure-associated infraslow oscillations throughout the brain. Our data showed that infraslow shifts propagate both within the same brain area and across different brain areas (Fig. 6 D, E, F). The propagation of seizure-associated SD waves throughout the brain was analyzed for a cohort of 9 seizures across 5 patients. To measure SD spread, we regressed the location of each recording electrode against the timing of its infraslow minima during and after the seizure interval (where *t*= 0 is when the first channel experiences an infraslow minimum, Fig. 6G). The correlation of infraslow minima delay to relative distance from the first observed infraslow minima was found to be significantly positive in 3 out of 9 individual seizures, as well as across the multi-patient cohort, providing evidence of SD spread throughout human brain tissue during and after seizures (Fig. 6H; *F*(1, 6.746)*=* 3.024; *p=* 0.027; see Supplemental Fig. 3). Additionally, the propagation velocities were determined with the best-fit line slopes of minima timing and location for each seizure. These calculated propagation velocities ranged from 7.57 to 111.19 mm/min with a median of 25.92 mm/min (Fig 6I). This collective data demonstrates that these large, infralow oscillations propagate in the brain, show diverse effects on ictal activity, and are very likely SDs.

## Discussion

Our findings indicate that SDs significantly influence ictal activity and play a role in seizure shortening or termination. Additionally, propagating SDs are a distinct feature of seizures in patients with intractable epilepsy, potentially affecting seizure initiation, spread, and termination. This highlights the need for increased attention and consideration of SDs when monitoring epilepsy patients. We argue that the insights provided from this study enhance our understanding of the mechanisms behind spontaneous seizure termination and could inform strategies to help terminate life-threatening seizures, such as status epilepticus.

In our preclinical model of seizures, we found that 33% of seizure-like events (SLEs) were associated with a SD at ictal termination, and these SLEs were significantly shorter than those that did not end with a SD. This data provides evidence that SDs can be a trigger for seizure termination and could be a potential mechanism that curtails the duration of seizures. Another key result from our study is that SDs do not appear to be driven by or correlate with changes in the [K^+^]_o_ levels created by the ictal discharges. This finding calls into question the relevance of changes in [K^+^]_o_ on SD induction during or following seizures. This leaves open an important question regarding how SDs might be induced during or following ictal events. While the potassium hypothesis of SD induction with seizures^38,58–60^ has been the most prominent for years, it has also been claimed that large increases in extracellular glutamate or simply strong neuronal depolarization can be key triggers as well.^21,60–63^ Of course, all these changes occur during ictal discharges, and it is possible that the exact right recipe of all three (strong neuronal depolarization combined with elevated glutamate and [K^+^]_o_) can trigger SDs. However, none of these potential mechanisms have been clearly established in SD induction following seizures.

Some recent work, in particular, provides evidence that contradicts the need for elevated [K^+^]_o_ or neuronal depolarization for SD initiation, where SDs were reliably triggered solely with the optogenetic chloride pump halorhodopsin.^64^ In these studies, it was confirmed that SD induction could occur during periods of low [K^+^]_o_ and when the neurons were hyperpolarized. It was also later confirmed in a follow-up study that SDs could not be triggered by hyperpolarization alone when using the optogenetic proton pump archaerhodopsin,^65^ suggesting elevated intracellular chloride as a potential mechanism for SD induction when triggered using halorhodopsin. This work suggests an alternative potential mechanism for SD induction during seizures because during ictal events, there is strong concurrent glutamatergic and GABAergic drive,^66–68^ providing a perfect storm for chloride loading of neurons. Indeed, raised intracellular chloride has been proposed to be a key mechanism behind the failure of strong feedforward inhibition that underlies seizure spread.^69–72^ It is therefore possible that chloride loading into neurons during ictal discharges causes SDs when chloride reaches a critical level, as with halorhodopsin-mediated chloride loading. Understanding how this process might contribute to the induction of a SD during seizures should be further investigated.

As reported by others,^14^ our preclinical model demonstrated significant cortical silencing following ictal events that ended in SDs compared to those that did not, along with a significant delay in subsequent ictal activity following SDs. Our work, along with others,^14^ suggests that SDs can be anti-ictogenic, preventing or delaying subsequent seizure activity. While SDs appear to curtail seizure activity in many instances, further work should investigate if the SDs increase morbidity when they associate with seizures. We further report an increase in low gamma activity during the ictal event present during the SLEs that end in SDs, which may suggest altered neuronal dynamics during a seizure that promotes the induction of or possibly signals the presence of a migrating SD. While this finding in our study of increased gamma is not clearly understood, increased gamma activity has been reported to be a hallmark of synchronous inhibitory neuronal activity.^73,74^ It could therefore create an interesting positive feedback mechanism that promotes SD induction due to the changes in intracellular chloride described above. It is also possible that a migrating SD triggered distally to the recording site could enhance feed-forward inhibition due to gap junction coupling between the inhibitory interneurons,^75^ resulting in increased gamma activity prior to the SD innervating and terminating the ictal event.

While our preclinical data suggest that SDs occur about 33% of the time with SLEs at ictal termination, our clinical data demonstrate that SDs may be far more frequent during seizures in human patients with intractable epilepsy. In our subset of human patients, all seizures analyzed were associated with a SD in some brain region during the ictal event (see Table 1 and Supplemental Fig. 2). This data contrasts with a recent study suggesting SDs mostly occur in the hippocampus during focal human seizures, a phenomenon the authors report to be an important mechanism to post-ictal wandering.^50^ In the human patients, the relationship between seizures and SDs appears complex, as in some situations SDs strongly associate with the end of the seizures, while other times the SD appears to “ride” on top of the spreading seizure wave (or the seizure rides the SD wave). This interplay between seizures and SDs has been documented in a rodent *in vivo* model of epilepsy,^76^ but how these two phenomena can in some cases co-occur in tandem is hard to understand given the general consequence of SD is strong cortical silencing and the general inability of neurons to fire action potentials for many minutes. A recent study could shed some light on this conundrum, where they demonstrate that depending on the level of complete SD innervation to a brain region, it can cause diverse effects from silencing when it completely innervates tissue to excitation when the innervation is limited.^77^ Therefore, in our study, instances where seizures and SD appeared to co-occur might be due to the SD not fully innervating the recording site, leading to the appearance of both phenomena simultaneously. The importance of this finding to how seizures and SDs interplay with human epilepsy will require further investigation.

These results highlight that SDs are not only a hallmark of experimental seizures but are also a critical part of human seizures in patients with epilepsy. In fact, our study suggests that SDs are likely a more prominent occurrence with seizures in surgical epilepsy patients than we see with experimental models, indicating we have been undervaluing the importance and implications of SDs during seizures. How this dynamic interplay between seizures and SDs shapes long-term seizure prevalence and morbidity remains to be determined but clearly warrants persistent and careful study, as do the mechanisms that lead to SD induction. Whether SD induction could be used to curtail life-threatening seizures is also worthy of investigation, even in light of the proposed contribution of SDs to SUDEP.^27,28^ The evidence presented herein suggests SDs are not uncommon during seizures and while in some cases they might be able to migrate to the brainstem, as seizures can,^78,79^ this is not likely the case in many situations and in some cases, seizures like status epilepticus could be more dangerous to the patient than the occurrence of a SD. This current study provides critical groundwork for such investigations.

## Acknowledgements

The authors thank Jyun-you Liou and Cameron Metcalf for their feedback on this work. The authors would also like to thank Isaac Stubbs for support in helping generate figures for this manuscript.

## Funding

The Brain Research Foundation, Brigham Young University Colleges of Life Sciences, Medical Research Council (UK).

## Competing Interests

The authors report no competing interests.

## Availability of data

The data and code that support the findings of this study are available from the corresponding author upon request.

---

Table 1 Seizure and SD Incidence Across Brain Area

## Notes

### Competing Interest Statement

The authors have declared no competing interest.

### Summary of Updates

The manuscript has been updated to refelct the correct author list and affiliations.

