## Supplementary figures and images for "Human and Rodent Seizures Demonstrate a Dynamic Interplay with Spreading Depolarizations"

### Supplemental Fig. 1

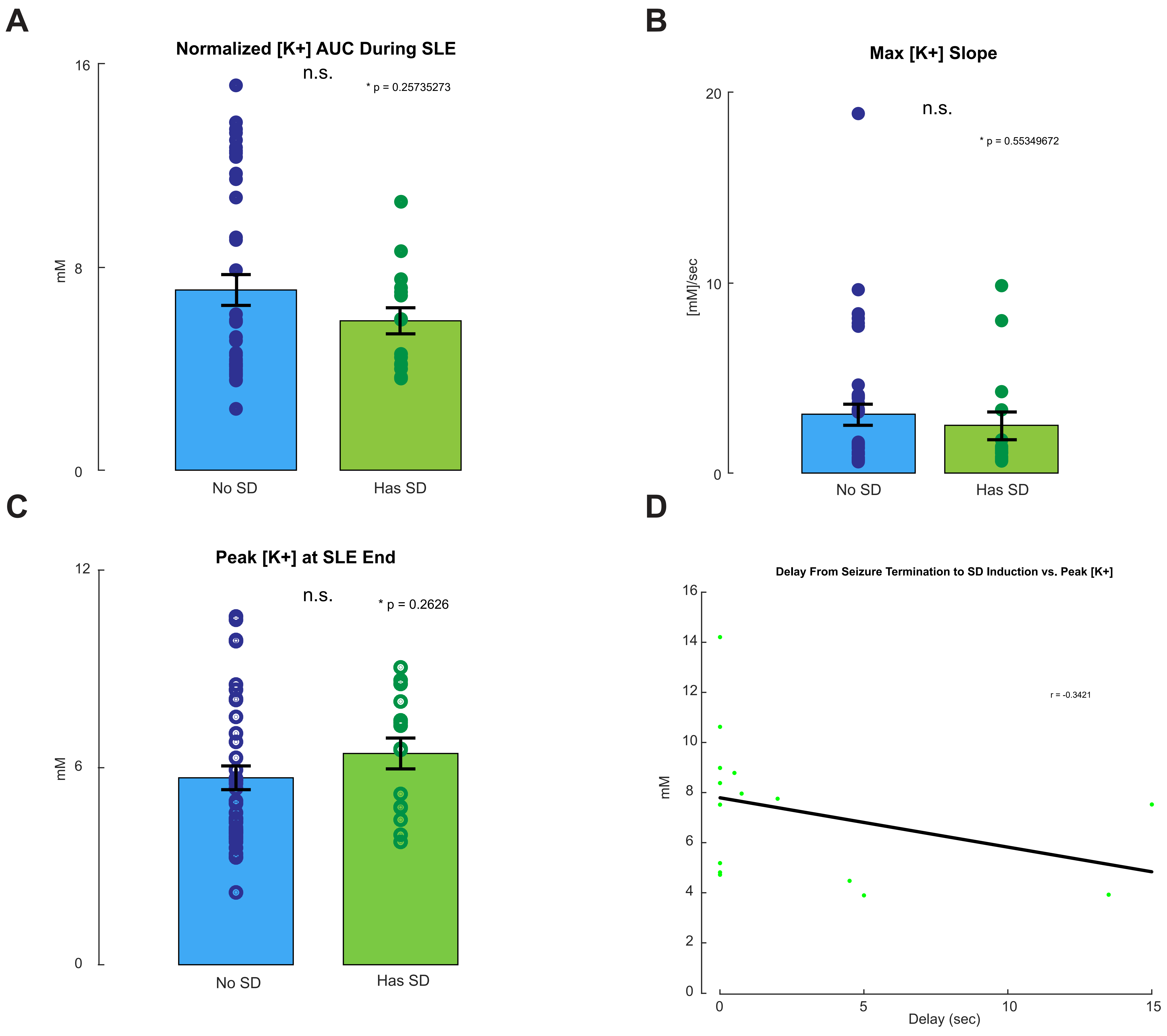

### Supplemental Fig. 2

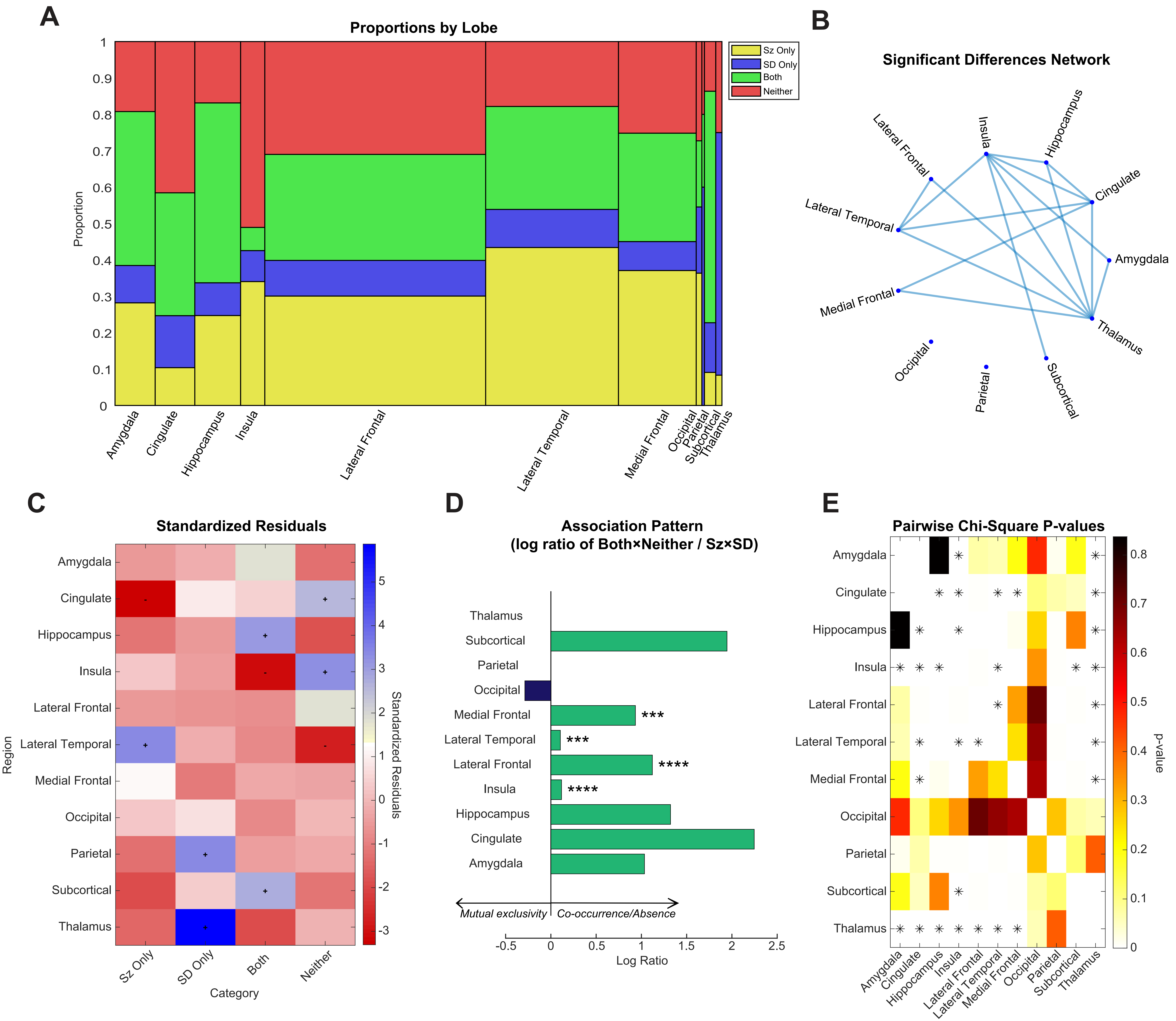

### Supplemental Fig. 3

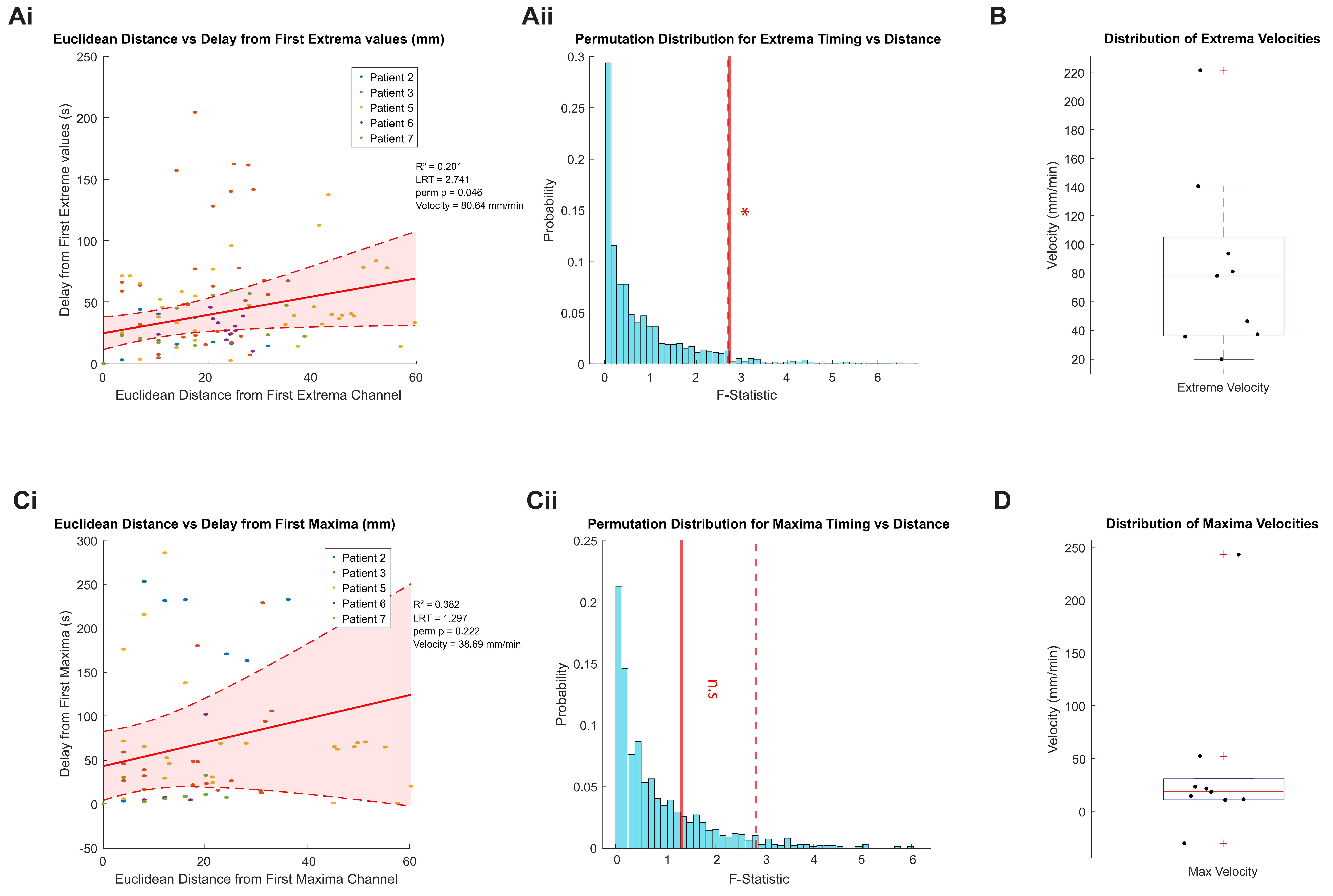
