## Supplemental Figure Legends for "Human and Rodent Seizures Demonstrate a Dynamic Interplay with Spreading Depolarizations"

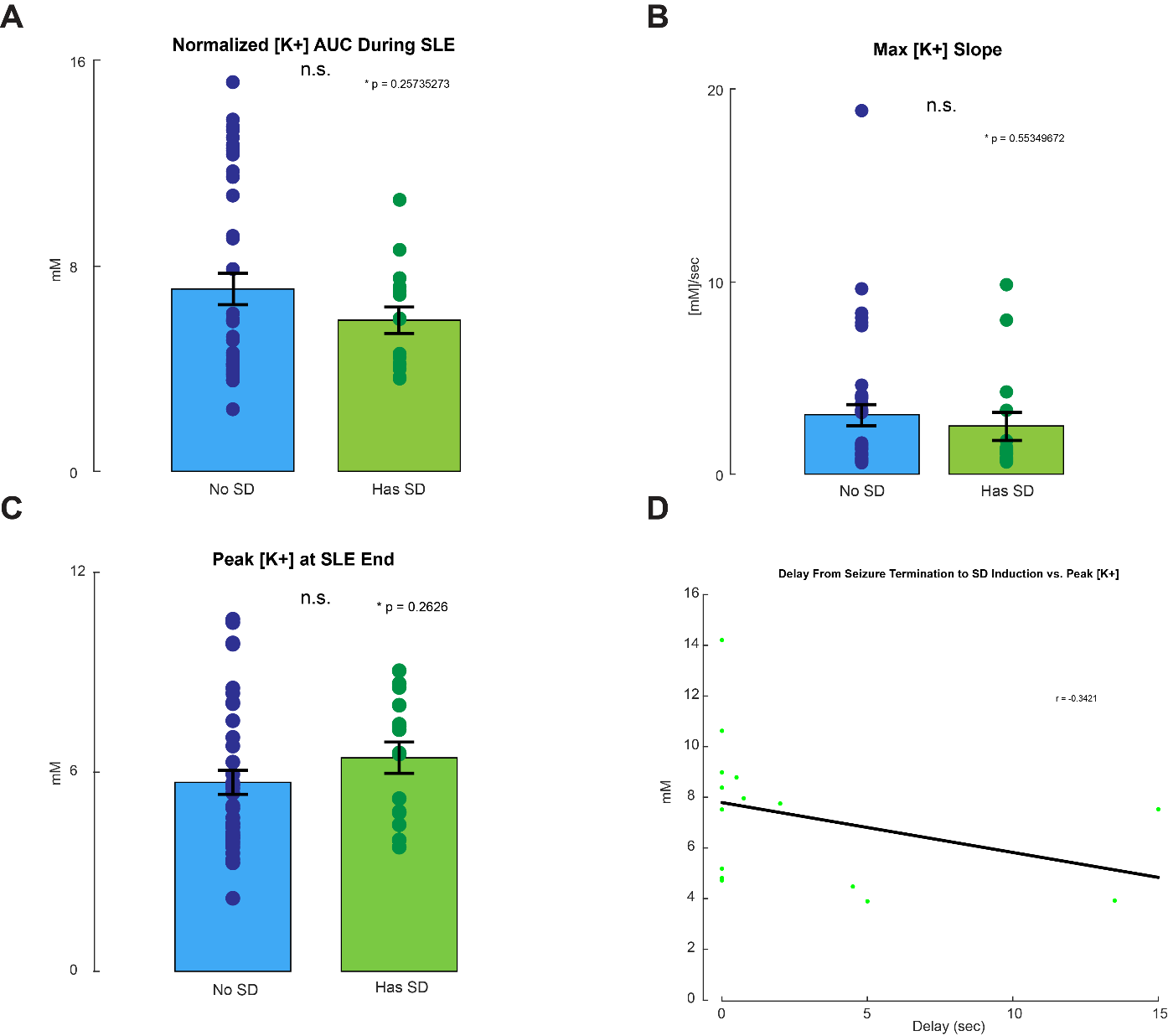

**Supplemental Figure 1 – Additional extracellular [K^+^] measures do not correlate with the presentation of SDs during or following SLEs (A)** The normalized [K^+^] area under the curve during SLEs did not show a significantly different [K^+^] load per unit time between SLEs that did terminate in a SD and those that did not (SLEs w/o SD, *n*= 41; SLEs with a SD, *n*= 15; two-tailed t-test, *p*= 0.257). **(B)** No significant difference was found between the maximum magnitudes of the [K^+^] change during the SLEs that did terminate in a SD and those that did not (SLEs w/o SD, *n*= 41; SLEs with a SD, *n*= 15; two-tailed t-test, *p*= 0.553). **(C)** Peak [K^+^] at the end of SLEs was not found to be significantly different between SLEs terminating in a SD and those that do not (SLEs w/o SD, *n*= 41; SLEs with a SD, *n*= 15; two-tailed t-test, *p*=0.263). **(D)** The peak [K^+^] reached during the SLE did not show a strong correlation to the time delay from the SLE termination to the SD induction (*r*= -0.342). All error bars represent SEM.

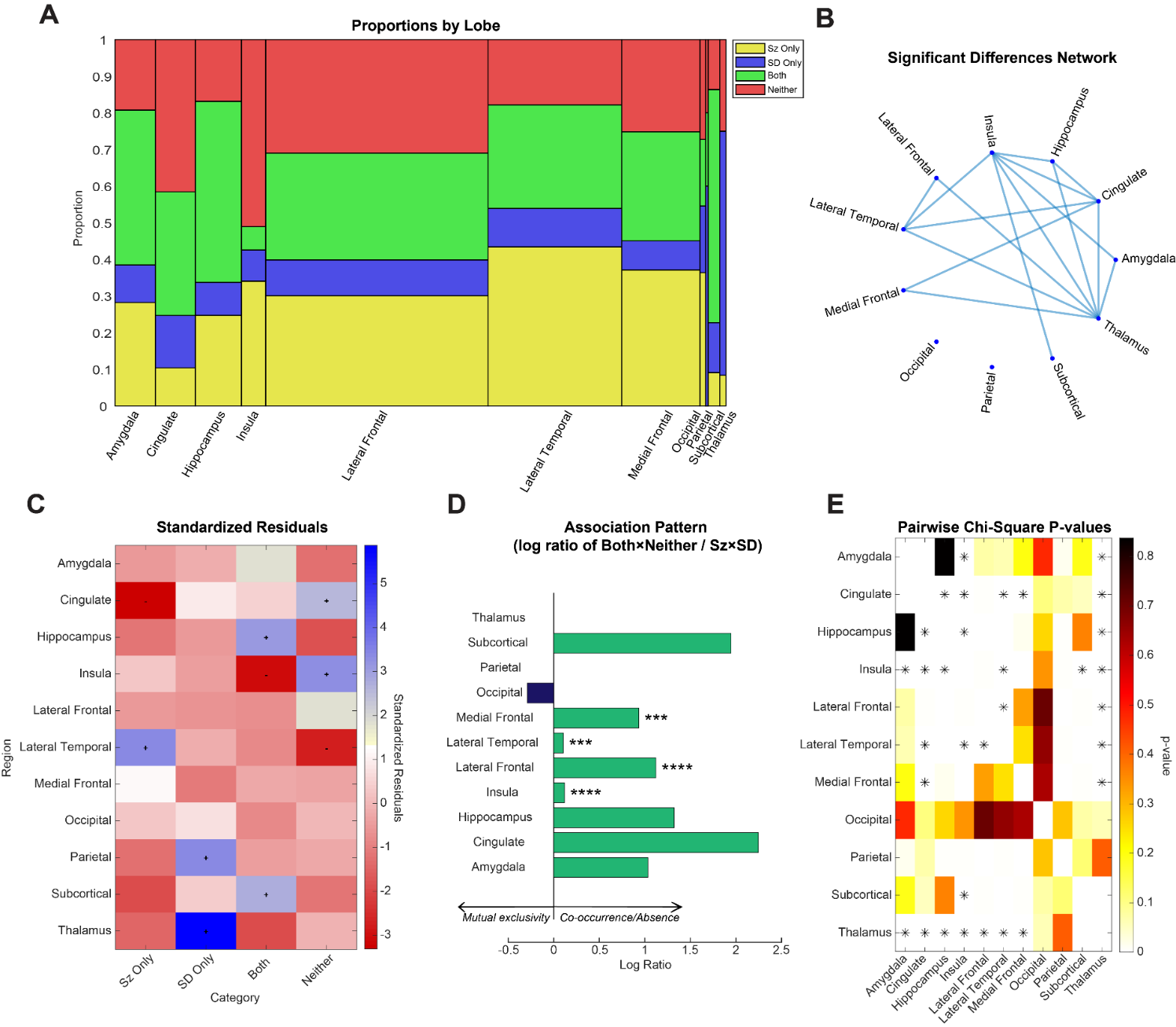

**Supplemental Figure 2 – Incidence of seizures and SDs significantly differ across brain areas in human patients (A)** Quantification of the distribution of observation category for each brain area. The width of each bar is proportional to the counts out of the total data set. **(B)** Network graph showing the brain areas with significantly different distributions of seizure and SD occurrence. A line indicates the pair is significantly different (pairwise post-hoc chi-square tests with Bonferroni correction for multiple comparisons). **(C)** Summary of the standardized residuals for each brain area. A ‘+’ or ‘-‘ indicates significant tendency for preference or aversion to the category, respectively (chi-squared residuals test; Bonferroni correction for multiple comparisons). **(D)** The medial frontal, lateral temporal, lateral frontal, and insula brain areas showed a significant preference for co-occurrence of seizures and SDs (G-test for association, ***, *p*< 0.001; ****, *p*< 0.0001). **(E)** Summary of the network graph shown in **B** (pairwise post-hoc chi-square test with Bonferroni correction for multiple comparisons). Significance less than the Bonferroni adjusted *p*-value is indicated by *.

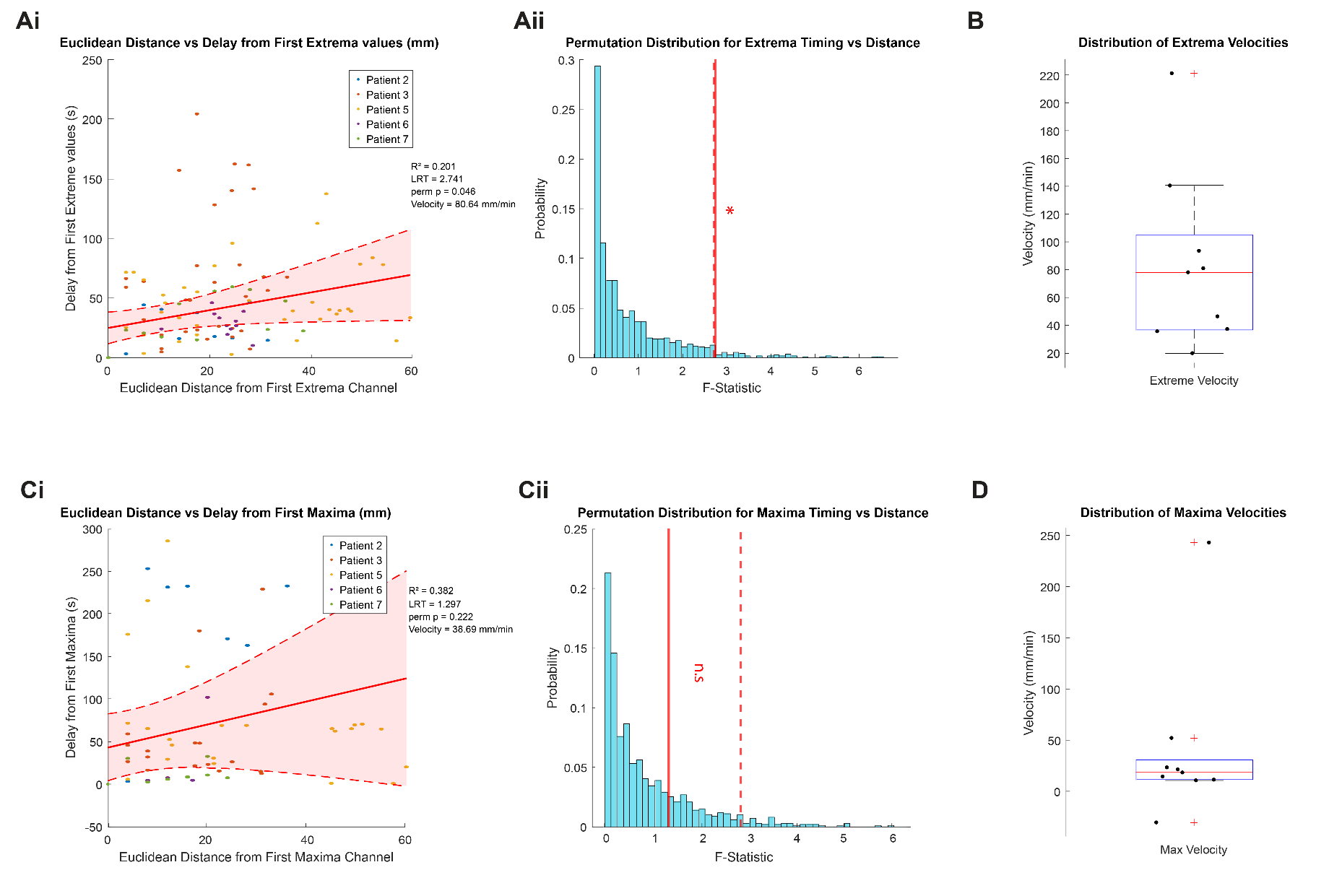

**Supplemental Figure 3 – Significant propagation of SD waves is seen based on most extreme values but not when based on maxima (A**) Combined propagation of SD waves was significant when propagating SD waves were identified by their most extreme values. **[Ai]** Correlation of electrodes showing temporal delay (vertical axis) and distance from the first DC shift minima (of each seizure recording) to display a DC shift that co-occurs with a seizure. The solid line represents the line of best fit and the dashed lines the 95% confidence interval (*n*= 108 electrodes from 9 seizures, *r*= 0.201, velocity= 80.64 mm/min). **[Aii]** The permutation distribution plot of the ANOVA F-statistic (using mixed effects model) over 10000 permutations shuffling the extrema DC shift delays and electrode locations (*p*= 0.046). **(B)** Distribution of the SD velocities for the 9 seizures from **A**. **(C**) Combined propagation of SD waves was significant when propagating SD waves were identified by their extrema of greatest magnitude. **[Ci]** Correlation of electrodes showing temporal delay (vertical axis) and distance from the first DC shift minima (of each seizure recording) to display a DC shift that co-occurs with a seizure. The solid line represents the line of best fit and the dashed lines the 95% confidence interval (*n*= 70 electrodes from 9 seizures, *r*= 0.382, velocity= 38.69 mm/min). **[Cii]** The permutation distribution plot of the ANOVA F-statistic (derived from the mixed effects model) over 10000 permutations shuffling of the maximum DC shift delays and electrode locations (*p*= 0.222). **(D)** Distribution of the SD velocities for the 9 seizures from **C**.

**Supplemental Video 1 – Calcium imaging and electrophysiology of SLE and SD**

**Supplemental Video 2 – Human GUI visualization of a seizure and SDs**
